# Effect of vessel compression on blood flow in microvascular networks: implications for tumour tissue hypoxia

**DOI:** 10.1101/2023.04.06.535833

**Authors:** Romain Enjalbert, Timm Krüger, Miguel O. Bernabeu

**Author notes:** Equally contributing senior authors (Timm Krüger, Miguel O. Bernabeu).

## Abstract

The tumour microenvironment is abnormal and one of its consequences is that blood vessels are compressed. Vessel compression correlates with reduced survival rates, while decompression of vessels improves tissue oxygenation as well as increases survival rates. Vessel compression contributes, at a single vascular bifurcation, to the increase of heterogeneity of red blood cell (RBC) transport. However, the effect that vessel compression has at a network level is unknown. This work numerically investigates the effect of vessel compression on RBC transport in microvascular networks. The key findings are that vessel compression both reduces the average haematocrit, and increases haematocrit heterogeneity, in vessels in the network. The mechanisms for these changes in haematocrit distribution are unravelled, and a parameter sweep shows that networks with lower inlet haematocrits are more susceptible to haemodilution from vessel compression over a wide range of compressed fraction of a network. These findings provide a theoretical underpinning for the link between vessel compression and tumour tissue hypoxia.

## Introduction

Solid tumours have an abnormal microenvironment, leading to tumour tissue hypoxia [1]. Hypoxia is an undesirable trait as it is associated with reduced patient survival rates through two separate mechanisms: increased tumour aggressiveness and resistance to tumour treatment [2, 3]. More recent therapeutic avenues have not translated to expected patient benefit, and the presence of hypoxia is one of the causes for that [4–6]. Blood vessel compression is one specific tumour microenvironment abnormality [7–9]. Previous work correlates the degree of vessel compression with reduced survival rates [8] and shows that pharmacologically decompressing vessels improves survival rates [10]. Furthermore, vessel compression is associated with tissue hypoxia as well as increased oxygen heterogeneity in the tissue [10].

The cause for increased hypoxia and increased oxygen heterogeneity due to vessel compression has not been fully elucidated [10]. At microvascular bifurcations, red blood cells (RBCs) heterogeneously partition to the child branches due to the finite size of RBCs [11]. Our previous work has shown that another structural abnormality, reduced interbifucation distance, changes the partitioning of the RBCs to the child branches at individual bifurcations, leading to increased tissue oxygen at a network level [12]. We previously showed that, at a single vascular bifurcation, vessel compression leads to a more heterogeneous partitioning of RBCs in the child branches of a bifurcation [13]. Given RBCs’ role in the transport of oxygen in blood [14], it follows that oxygen transport is altered too. Previous work suggests that increased resistance to blood flow due to vessel compression leads to reduced perfusion to the blood vessels [15, 16]. However, how vessel compression affects haematocrit in blood vessels at a network level is unknown, nor has it been elucidated whether it drives tissue hypoxia.

In the current study, we find that vessel compression both reduces the average haematocrit, and increases haematocrit heterogeneity, in vessels in the network. We further show that networks with lower inlet haematocrits are more susceptible to haemodilution from vessel compression. These findings provide a theoretical underpinning for the link between vessel compression and tumour tissue hypoxia.

## Methods

Blood is modelled in two different ways. The first method is more accurate at a high computational cost, whereas the other method is based on a more efficient reduced-order model. The former, more highly resolved simulations serve to inform the latter reduced-order model.

### Particulate blood flow

With the first method, we treat blood as a suspension of deformable RBCs in a continuous plasma phase. The plasma is assumed to be a continuous Newtonian fluid, and the viscoelastic properties of plasma are not a leading-order effect in the regime investigated [17]. The non-Newtonian behaviour of blood arises from the presence of the deformable RBCs which are modelled as hyperelastic membranes.

The physics of the particulate blood flow is characterised by the Reynolds number and the capillary number. The Reynolds number quantifies the ratio of inertial to viscous forces in fluid flow. The capillary number quantifies the deformability of the particles, where higher capillary numbers correspond to softer particles. The Reynolds and capillary numbers are defined as

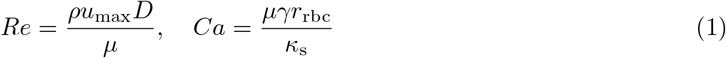

where *ρ* is the fluid density, *u*_max_ is the maximum velocity in the channel, *D* is the channel diameter, *μ* is the dynamic viscosity of the suspending fluid, *γ* is a typical shear rate in the system, *r*_rbc_ is the rest radius of the RBC, and *κ*_s_ is the elastic modulus of the RBC. In our simulations, the Reynolds and capillary numbers are set to 1 and 0.1 respectively, see [13] for more information.

The algorithm for solving for particulate blood flow is implemented in the open-source software HemeLB (https://github.com/hemelb-codes/hemelb) [18]. Additional information on the model and the numerical scheme can be found in the supplementary material and the references therein.

### Network blood flow

With the second method, blood flow is treated in a one-dimensional network model imposing Poiseuille’s law on each vessel segment, with additional components to the model accounting for the Fahraeus effect, the Fahraeus-Lindqvist effect, and partitioning of RBCs at bifurcations [19]. Additional information on the model and numerical scheme can be found in the supplementary material and the references therein. When vessels are treated as compressed, the effect of abnormal partitioning of RBCs and increased resistance can be numerically isolated, as they are separate steps in the numerical scheme. Separating the two effects allows us to study the effect of increased resistance without abnormal partitioning, called increased resistance case (IR), and the effect of abnormal partitioning of RBCs without increased resistance, called abnormal partitioning case (AP). This approach leads to four different types of simulation cases, shown in Table 1.

**Table 1.**
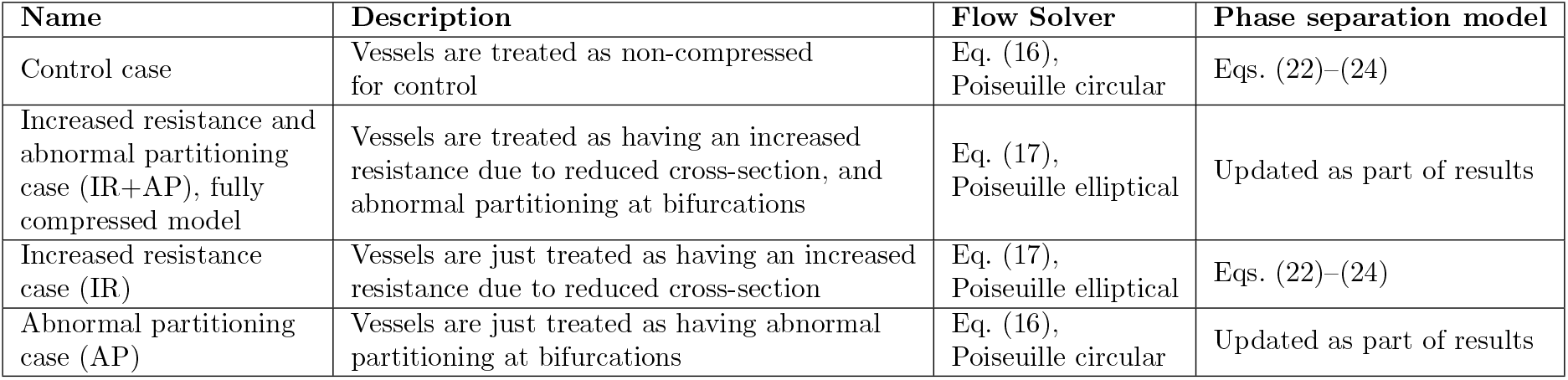
Four possible scenarios for how vessels are treated in network simulations, which determines how the compressed vessels, in red in Fig. S1, are treated.

| Name | Description | Flow Solver | Phase separation model |
| --- | --- | --- | --- |
| Control case | Vessels are treated as non-compressed for control | Eq. (16), Poiseuille circular | Eqs. (22)–(24) |
| Increased resistance and abnormal partitioning case (IR+AP), fully compressed model | Vessels are treated as having an increased resistance due to reduced cross-section, and abnormal partitioning at bifurcations | Eq. (17), Poiseuille elliptical | Updated as part of results |
| Increased resistance case (IR) | Vessels are just treated as having an increased resistance due to reduced cross-section | Eq. (17), Poiseuille elliptical | Eqs. (22)–(24) |
| Abnormal partitioning case (AP) | Vessels are just treated as having abnormal partitioning at bifurcations | Eq. (16), Poiseuille circular | Updated as part of results |

### Artificial network generation

The networks are generated using the open-source software Tumorcode [20–22]. It uses an algorithm to randomly generate vessel networks on a mesh. The algorithm is based on Murray’s law and geometrical properties for capillaries, arterioles, and venules. Tumorcode also has the ability to reproduce the evolution of a tumour vessel network in the *in silico* generated vascular network. However, for the purpose of the research here only the healthy networks are used, to isolate the network from other structural abnormalities. An example of a generated network is depicted in Fig. S1. We use the two-dimensional default network generator, with a single inlet and a single outlet. The single inlet and outlet allow the setting of the boundary conditions as pressure values, where the absolute value does not have an effect on the haematocrit output of the simulations. The two-dimensional networks are generated within a square surface of 5000 *μ*m by 5000 *μ*m.

Within the network, we treat some vessels differently, detailed in Table 1, which we call the compressed vessels or compressed region. Initially, the compressed vessels are in the centre of the network and are vessels that have both ends within a 1000 *μ*m radius of the centre of the vessel network. We also vary the fraction of compressed vessels, using a radial model and a random model. The radial model is a model for tumours where the solid stress is higher in the centre, thereby compressing vessels more in the centre of the tumour [23]. In the radial model, the compressed vessels are the vessels closest to the centre of the network (see Fig. S1a–f). The random model is an alternative model where compressed vessels are not deterministically situated. In the random model, a random selection of vessels is compressed to reach the desired fraction of total vessels (see Fig. S1g–l).

### Processing results

To process the results, we also define a set of critical bifurcations. A critical bifurcation is defined as a diverging bifurcation where

1. one of the child branches has every possible path emanating from it going through the compressed region and
2. the other child branch has at least one path emanating from it not going through the compressed region.

Fig. S2 illustrates one of these critical bifurcations.

In addition, to quantify the effect that vessel compression has on the reduction of haematocrit in the networks, we define a metric which is a proxy to quantify haematocrit reduction in a network. The metric uses the basic form

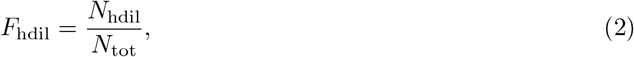

where *F*_hdil_ is the fraction of haemodiluted vessels in the network, *N*_hdil_ is the number of haemodiluted vessels in the network, and *N*_tot_ is the total number of vessels in the network. The condition for haemodilution is that the haematocrit in a given vessel has a relative value of 50% or less of the haematocrit of the same vessel in the control case.

## Results

### Adapting the reduced-order model for compressed vessels

To model the partitioning of RBCs in compressed vessels at a network level, we adapt an existing reduced-order model, developed by Pries et al. [19, 24], to account for vessel compression. Fig. 1a shows a flow chart illustrating the method we used to update the reduced-order model. We choose to adapt Pries’ model due to its robustness and ubiquity in the literature [24].

**Figure 1.**
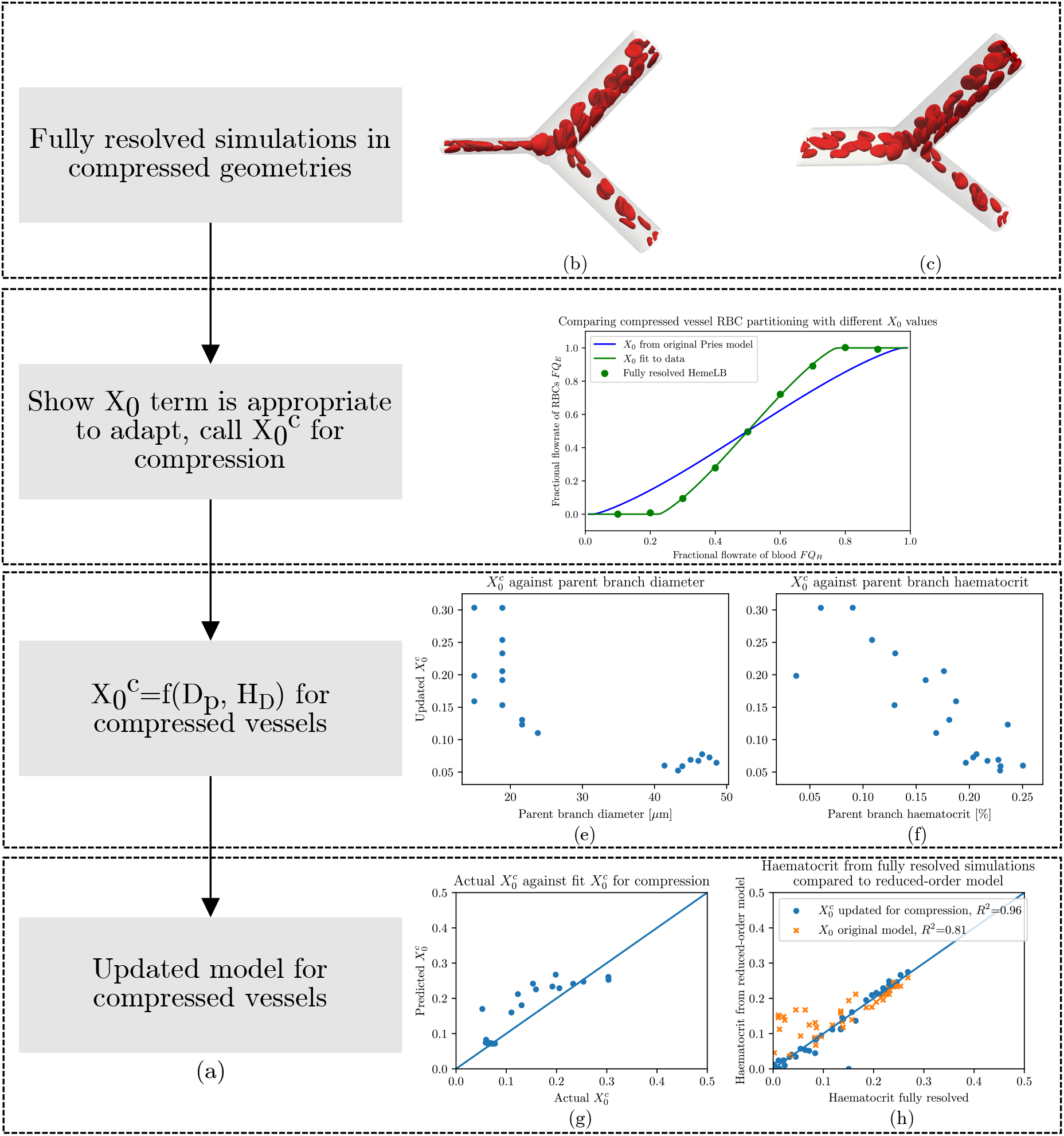
(a) Flow chart of the process to calculate the values for 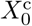 for RBC partitioning in compressed vessels. (b) Snapshot of a fully resolved cellular blood flow simulation. (c) Rotated snapshot of same simulation to show compression has an elliptical cross-section. (d) Plasma skimming curve. Green points are from varying the flow ratio in a bifurcation geometry of 33 *μm* with a compression before the bifurcation and an inlet haematocrit of 20%. Compares how well the original phase separation empirical model works (blue line) and how changing solely *X*_0_ improves the fit (green line). *X*_0_ for the green line is obtained through a fit to the data using the non-linear least-squares method. (e) 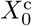 calculated from fully resolved simulations with Eq. (3) against the parent branch diameter. (f) 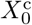 calculated from fully resolved simulations with Eq. (3) against the parent branch haematocrit. (g) Predicted 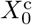 from Eq. (6) against actual 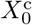 as calculated from fully resolved simulations. (h) Predicted haematocrit (using updated 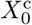 in blue and original *X*_0_ in orange) against the fully resolved haematocrit. The diagonal line in (g) and (h) is a visual aid to see how close the predicted values are to the values obtained from fully resolved simulations.

We initially demonstrate that the term that needs adapting in the model is *X*_0_ (see the supplementary material for more details on the model). We run a series of resolved RBC simulations (see Fig. 1b–c), varying the flow ratio in a bifurcation geometry with 33 *μ*m branches with a compression before the bifurcation (the parent branch is compressed with an aspect ratio of 4.26, as per [13]) and an inlet haematocrit of 20%. We plot the results as a plasma skimming curve, Fig. 1d, and show that by adapting *X*_0_ we obtain a close fit to the data generated in compressed vessels.

Supported by this evidence, we set out to find a new functional form for *X*_0_ for cases when vessels are compressed. We fully resolve a set of 20 bifurcations with HemeLB simulations (see Table S3). The flow ratios, haematocrit values, and diameters in Table S3 were taken from the diverging bifurcations in the network that will be used (see Fig. 2a) to be representative of the bifurcations that the updated phase separation model will resolve. The haematocrit and flow ratio in these bifurcations were obtained by solving the Poiseuille flow through the entire network using the standard phase separation model for RBC partitioning [19, 24].

**Figure 2.**
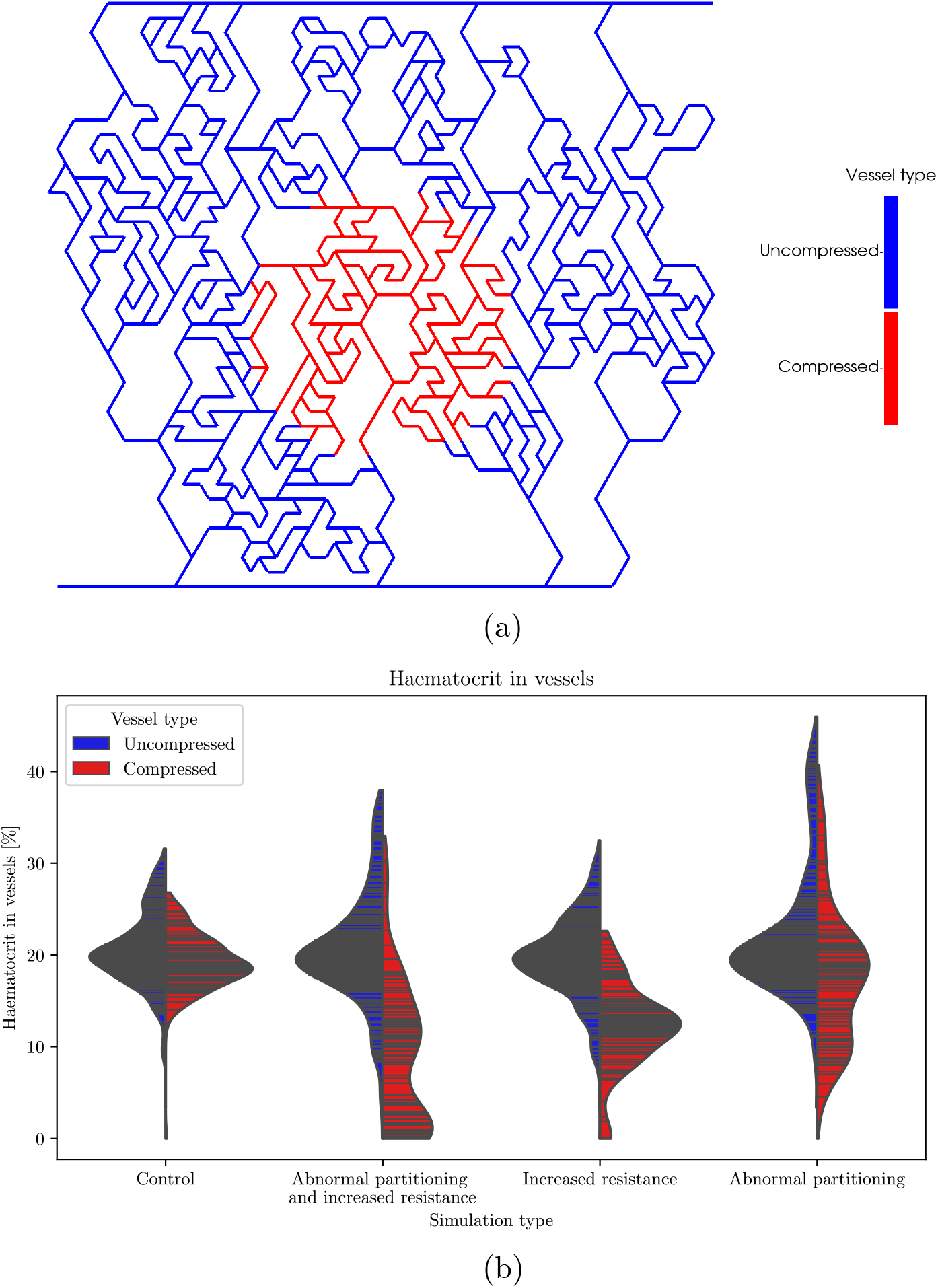
(a) Depiction of the network in which the blood flow simulations are performed. The blue vessels are treated as normal (uncompressed) blood vessels, the red vessels are treated as compressed vessels, also called compressed region. The inlet is at the bottom left, and the outlet is at the top right. (b) Violin plots of haematocrit distribution within the vessel network. Blue and red lines correspond to blue and red vessels in (a), respectively. The ‘control’ case treats all vessels as normal vessels. The ‘abnormal partitioning and resistance’ case treats compressed vessels as having an increased resistance and abnormal partitioning. The ‘abnormal partitioning’ case treats compressed vessels as having just abnormal partitioning. The ‘resistance’ case treats compressed vessels as having just increased resistance. See Table 1 for further details about the four cases.

Next, we need to find the values of *X*_0_ for the results of these simulations that match the child branch haematocrit values from the fully resolved simulations. We analytically invert the main equation of the Pries model to make *X*_0_ the subject of the equation,

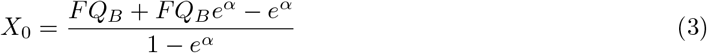

where *FQ*_*B*_ is the fraction of blood flowing to a child branch and *α* is defined as

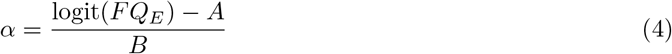

where *FQ*_*E*_ is the fraction of RBC flowrate flowing to a child branch and *A* and *B* are terms defined in the original empirical model [19, 24].

We calculate the updated value of *X*_0_ for compressed vessels, 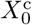, from Eq. (3) and *FQ*_*E*_ values from Table S3. Fig. 1e–f show that 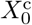 is inversely proportional to *D* and decreases linearly with *H*. Therefore, we keep the original functional form, and update the pre-factor:

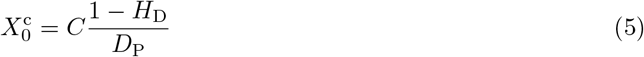

where *H*_D_ is the discharge haematocrit in the parent branch, *D*_P_ is the diameter of the parent branch, and *C* is the pre-factor to be determined for the new functional form for 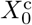. We use a non-linear least-squares method to fit *C* and obtain a value of 4.16, yielding

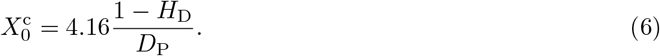

We now verify that the updated form of 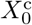 accurately captures the RBC partitioning from the fully resolved simulations. Fig. 1g shows that Eq. (6) does not perfectly capture the analytically calculated value of 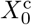. However, Fig. 1h shows that the haematocrit in the child branches, the desired output of the model, is well predicted by the updated term 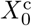 with an *R*^2^ value of 0.96, and fits the data much better than the original model, with an *R*^2^ value of 0.81.

### Vessel compression reduces haematocrit and increases haematocrit heterogeneity

We start by investigating how the compressed vessels alter the distribution of RBCs, and therefore haematocrit, at a network level. We run the network model for blood flow in an artificially generated network, shown in Fig. 2a where we treat the vessels in red as being compressed according to the description in Table 1, which for the control case implies all the vessels in the network are treated as uncompressed.

Fig. 2b shows that, when the vessels are treated with the fully compressed model, there is a reduction in the average haematocrit of the compressed vessels compared to the control from 19.2% to 8.8%. Fig. 2b also shows that the distribution of haematocrit within the compressed vessels is wider, ranging from [0.0%, 26.8%] in the control and increasing to [0.0%, 32.9%] in the fully compressed case which also becomes bimodal. The frequency of vessels with 0% haematocrit is increased from 1 vessel in the control case to 11 vessels in the fully compressed case.

Next, we separately investigate the effects of increased resistance and abnormal partitioning. When comparing the cases with increased resistance and the control cases, one initially sees that the distribution remains unimodal with a longer tail, Fig. 2b. Contrarily, the distribution of the case with abnormal partitioning is bimodal. This bimodal distribution indicates that the increased heterogeneity of haematocrit results from abnormal partitioning of RBCs, rather than from increased resistance.

Finally, one sees that both the cases with increased resistance and abnormal partitioning reduce the average haematocrit in the compressed region down to 12.1% and 17.7%, respectively, compared to 19.2% in the control. This finding suggests that there are two separate mechanisms that contribute to the reduction of average haematocrit in the network. We will investigate those mechanisms next.

### Mechanism that reduces haematocrit due to increased resistance

Next, we explain the mechanism through which increased flow resistance of the compressed vessels reduces the haematocrit within the compressed region. We hypothesise that this effect results from diverting flow away from the compressed vessels due to their higher resistance to flow, according to Poiseuille’s law. Therefore, as the phase separation effect disproportionally favours RBC flow into the higher-flowing child branch [11], RBC flow into the compressed region should be disproportionally reduced, thus decreasing the haematocrit in the compressed region.

To test our hypothesis, we investigate all 11 critical bifurcations in the network (see Methods and Fig. S2 for details). We compare the flow ratio at these critical bifurcations between the control case and the case with increased resistance. Fig. 3a shows that the flow ratio in the case with increased resistance is always reduced in the child branch which necessarily goes through the compressed region. This observation confirms that flow is diverted away from the compressed region.

**Figure 3.**
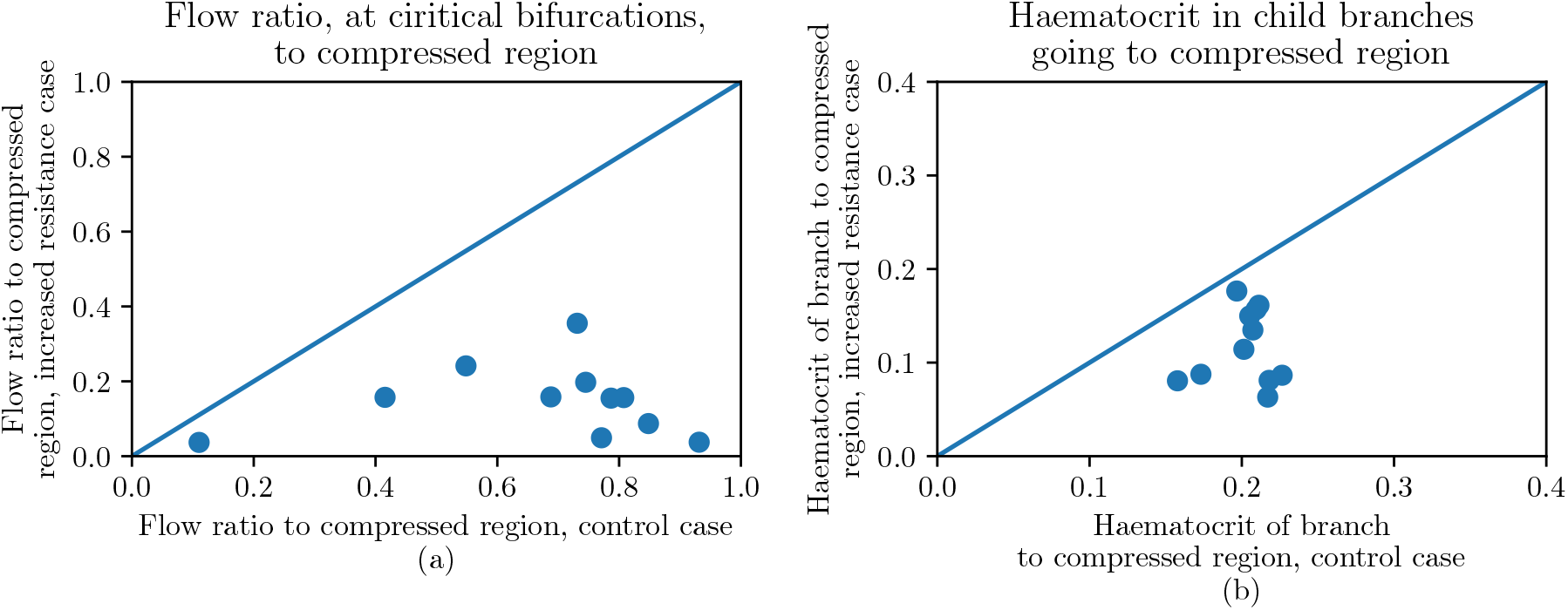
Effect of increased resistance at all 11 critical bifurcations in the vascular network (see Fig. 2 for network map). (a) Comparison of the flow ratio into the compressed region between the control case and the case with increased resistance, showing a reduced flow ratio into the compressed region when compressed vessels have an increased resistance. (b) Comparison of the haematocrit of the child branch going into the compressed region between the control case and the case with increased resistance, showing a reduced haematocrit in the branch flowing into the compressed region when there is an increased resistance. The diagonal line in (a) and (b) is a visual aid to see how close the values in the case of increased resistance are to the control case.

Next, we show how the decrease of flow into the compressed region leads to a reduction in haematocrit in the compressed region. We take the geometrical properties of the critical bifurcations and consider the haematocrit of the parent branch as obtained from the network simulations using the case with increased resistance. We then solve for the haematocrit distributions in the child branches of the critical bifurcations using the original phase separation model which contains the plasma skimming effect [11]. We prescribe the flow ratios in the child branches with and without the increased resistance and compare the resulting haematocrit distributions. Fig. 3b shows that the haematocrit of the vessel going into the compressed region is smaller for the case with increased resistance as compared to the control for all critical bifurcations, which confirms our hypothesis.

### Mechanism that reduces haematocrit due to abnormal RBC partitioning

As a next step, we investigate the effect that the abnormal partitioning has on haematocrit within the network. To this end, we use the model for abnormal partitioning which increases haematocrit heterogeneity in the child branches. We hypothesise that the reduced average haematocrit in the compressed region caused by abnormal partitioning alone is due to an enhanced network Fahraeus effect.

The network Fahraeus effect contributes to the reduction of average haematocrit at each successive bifurcation as the average haematocrit of the two child branches at a bifurcation is lower than the haematocrit of the parent branch [25].

To investigate the hypothesis, we first show that, at a vascular bifurcation where the higher flowing child branch is disproportionately enriched in haematocrit, the average haematocrit of the child branches is lower than that of the parent branch (branch 0):

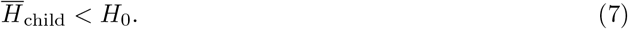

The supplementary material shows how to obtain this relation. We further define Δ*H*_1_ and Δ*H*_2_ as the differences in haematocrit between the parent branch and the two child branches (branches 1 and 2, where branch 1 is enriched):

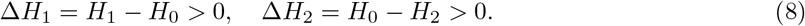

Fig. 4 shows the data from diverging bifurcations in the compressed portion of the network for the control case and the case with abnormal partitioning. It can be seen that Δ*H*_1_ and Δ*H*_2_ are larger in the case with abnormal partitioning compared to the control case. Further, the mean reduction in haematocrit, 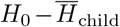, is increased in the case with abnormal partitioning compared to the control case. Thus, a stronger network Fahraeus effect occurs in the case with abnormal partitioning. Additionally, the enhanced heterogeneity in haematocrit due to the abnormal partitioning suffices to explain the wider and bimodal distribution of haematocrit within the compressed portion of the network observed in Fig. 2.

**Figure 4.**
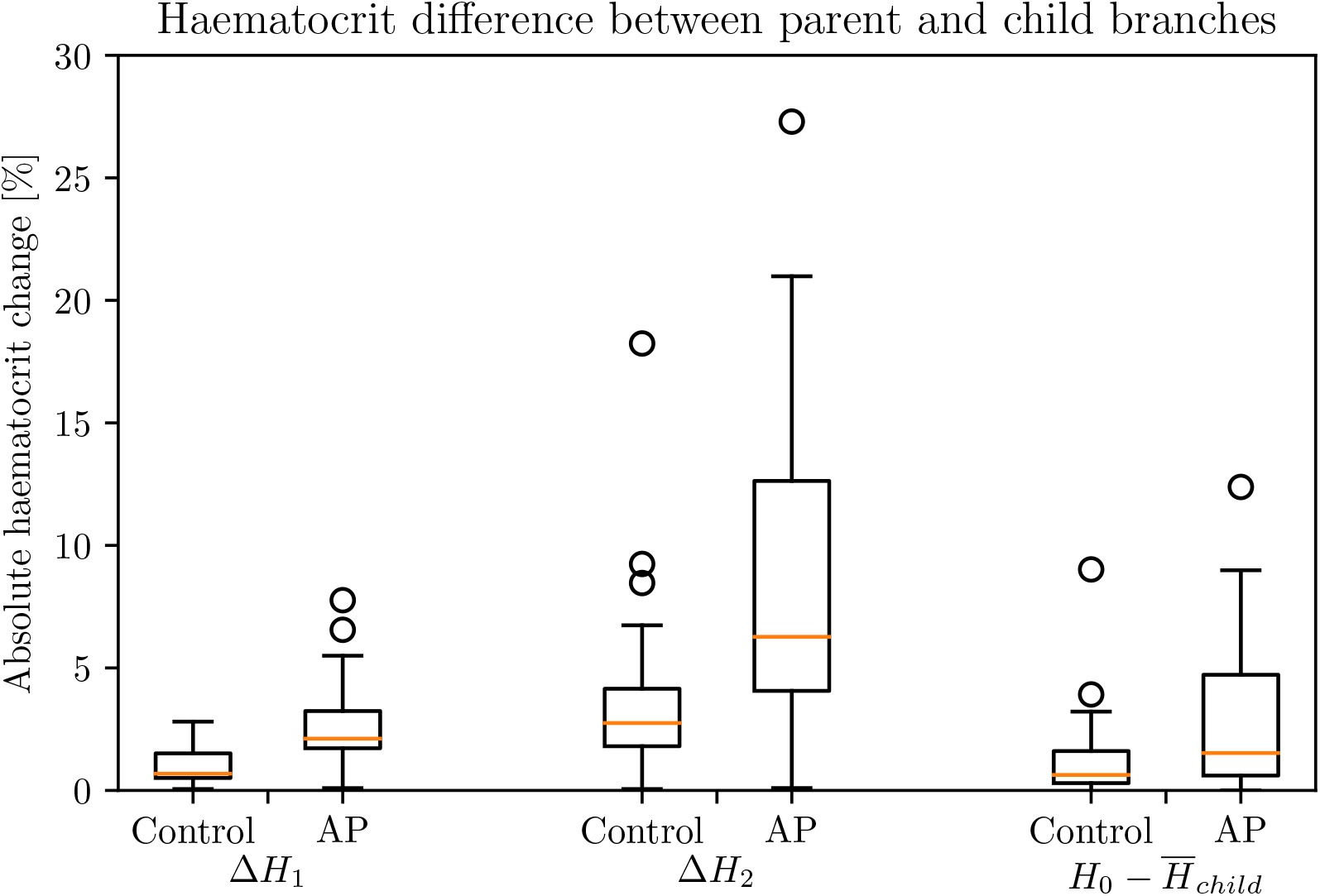
Left and middle: enrichment, Δ*H*_1_, and impoverishment, Δ*H*_2_, of the compressed vessels for the control case and the case with abnormal partitioning (AP). Right: reduction in average haematocrit between the parent branch and the average of the child branches, 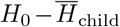, at a diverging bifurcation in the compressed region for the control case and the AP case.

### Network conditions for haemodilution

Finally, we investigate under what conditions networks are susceptible to haemodilution when vessel compression is present. We perform a parameter sweep over a wide range of inlet haematocrits to the network, from 2.5% to 30% in steps of 2.5%, and over the entire range of fraction of compressed vessels, using the fully compressed model (increased resistance and abnormal partitioning). As described in the Methods and in Fig. S1, we consider two models for the fraction of compressed vessels in a network, one where the more central vessels are compressed (radial model), and the other where random vessels are compressed (random model).

Fig. 5a shows the fraction of haemodiluted vessels in the radial model. The results show that haemodilution generally increases with the fraction of compressed vessels and with a decreasing inlet haematocrit. Fig. 5a also indicates that, although haemodilution is experienced over the entire parameter range, it is most pronounced when the inlet haematocrit to the network is relatively small, below 15%. In addition, this haemodilution effect is observed over a wide range of fraction of compressed vessels, suggesting that a low fraction of compressed vessels is sufficient for the network to experience haemodilution. Lastly, in the random model, Fig. 5b, it is interesting to see that the overall haemodilution trend is not very different compared to the radial model of compressed vessels, although the haemodilution is less pronounced.

**Figure 5.**
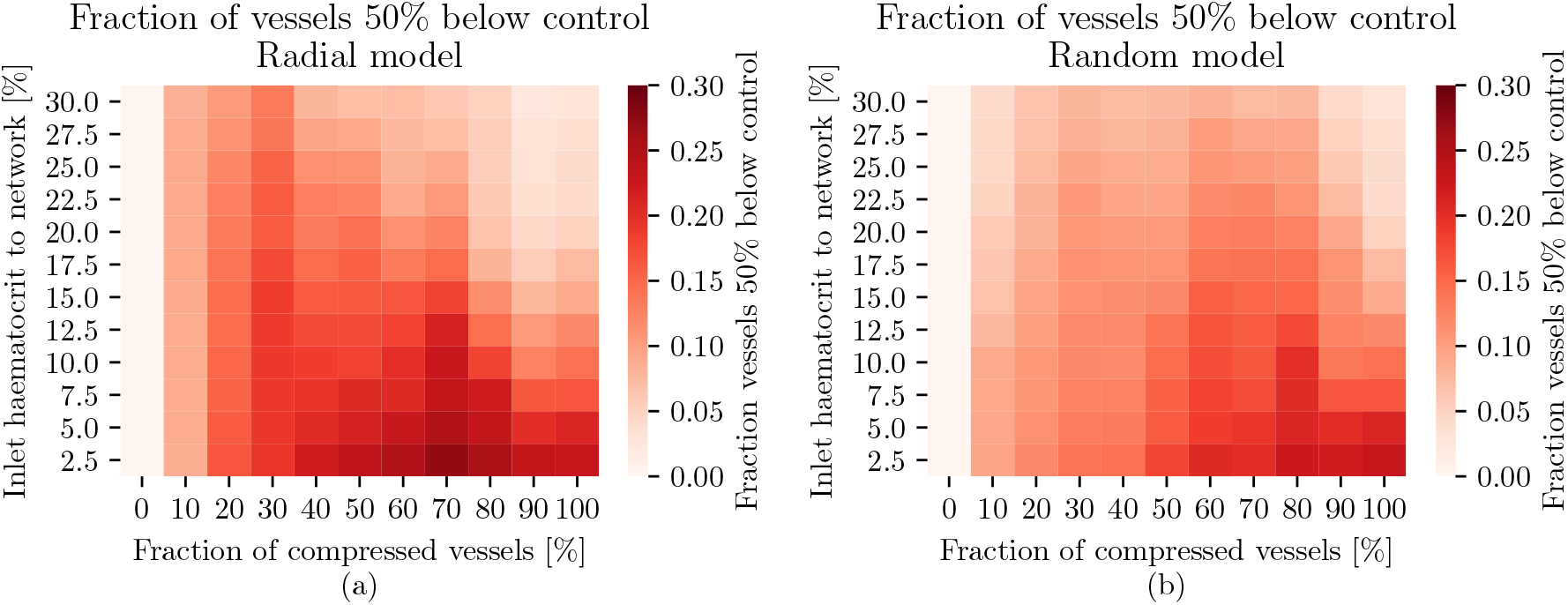
Heat map showing the fraction of haemodiluted vessels in the network for varying inlet haematocrit and fraction of compressed vessels in the network. Haemodiluted vessels are defined as vessels with a haematocrit of less than half of the haematocrit of the same vessel in the control case. (a) Radial model for the network, (b) random model for the network. See Fig. S1 for an illustration of the radial and random models.

We also observe from Fig. 5 that the highest fraction of haemodiluted vessels is not when the fraction of compressed vessels is highest. Indeed, beyond a fraction of compressed vessels of around 80%, haemodilution decreases again. We attribute this behaviour to the mechanism for haematocrit reduction in the networks. We previously identified that increased vessel resistance has the most pronounced effect on haematocrit reduction and that it depends on flow redirection. However, when a high fraction of the network is compressed, the opportunity for flow redirection is reduced, leading to a decrease in the effect of flow resistance within the network, thus reducing the effect of compression on haemodilution.

## Discussion

Vessel compression in solid tumours is associated with both tumour tissue hypoxia and reduced survival rates in patients. Conversely, pharmacological decompression of vessels has been shown to lead to an improvement in survival rates [10]. However, a complete description of the biophysical processes underpinning these associations, particularly in relation to oxygen transport, remains elusive. Gaining a mechanistic understanding into them may uncover novel therapeutic strategies [26]. In this study, we numerically investigate RBC distribution in partially compressed vascular networks, as a model of tumour blood flow. We start by deriving a reduced-order model for RBC partitioning at bifurcations in the presence of vessel compression. We next demonstrate that vessel compression both reduces average haematocrit and increases haematocrit heterogeneity throughout the network. We identify, for the first time, two mechanisms that synergise non-linearly leading to this effect. The first is increased vessel resistance, and therefore flow and haematocrit diversion, in the portion of the network undergoing compression; the second is abnormal RBC partitioning in the compressed bifurcations, which underpins haematocrit heterogeneity at a network level. Finally, we show that haematocrit reduction due to compressed vessels increases with reducing inlet haematocrit to the network and with increased fraction of compressed vessels, with a maximum at around 80% of compressed vessels.

There are several implications of the research in this work. Firstly, it provides a theoretical underpinning to explain reports from the literature, where animal models are shown to have higher tissue hypoxia and tissue oxygen heterogeneity when vessels are compressed [10], as well as the presence of plasma channels in tumour vessels [27]. We highlight the importance of RBC transport at the network level in the emergence of these phenomena. The results support the hypothesis from our previous work 13 stating that, in a few successive bifurcations, vessels can be depleted of RBCs as a consequence of vessel compression [13], leading to plasma channels [27].

Secondly, this work forms part of a larger corpus in the literature studying how tumour vascular abnormalities affect blood transport in tumour microvascular networks [12, 13, 28–31]. Previously, inter-bifurcation distance and increased vessel diameter have been studied in isolation [12, 13, 28–31], and this work does it, for the first time, for vessel compression. However, future work is necessary to identify the relative effect on blood flow, and tissue oxygenation, of the different structural abnormalities as well as their relevance *in vivo*. Identifying the most relevant structural abnormalities in tumour microvascular networks could open novel avenues for diagnosis and patient treatment planning [32]. Prognosis could be improved through phenotyping tumour vascular networks based on their structural abnormalities, as these correlate with survival rates [8]. Treatment planning could be improved through optimising treatment based on the present structural abnormalities, to improve treatment delivery or efficacy [33, 34].

Thirdly, the results support the notion, hypothesised in our previous work [13], that healthy vascular networks are protected against the effects of naturally occurring structural abnormalities for as long as their average haematocrit remains high. Diseased networks, however, are susceptible to a positive feedback loop whereby they are at a higher risk of haemodilution as they have lower average haematocrit than healthy networks [27].

## Acknowledgments

We acknowledge the contributions of the HemeLB development team. Supercomputing time on the ARCHER2 UK National Supercomputing Service (http://www.archer2.ac.uk) was provided by the ‘UK Consortium on Mesoscale Engineering Sciences (UKCOMES)’ under EPSRC grant no. EP/R029598/1, with computational support from the ‘Computational Science Centre for Research Communities (CoSeC)’ through UKCOMES. R.E. and M.O.B. gratefully acknowledge their funding as part of a Fondation Leducq Transatlantic Network of Excellence (17 CVD 03, https://www.mdc-berlin.de/leducq-attract) and EPSRC grant no. EP/X025705/1. R.E. is funded by The University of Edinburgh through a Chancellor’s Fellow PhD studentship.

## Supplementary material

### Particulate blood flow model

Below, details are given on the particulate blood flow model. The interested reader is referred to the literature for more details about the model and its numerical implementation [35, 36], which has been validated for the relevant effects studied within this work [37–39].

### Fluid model

The Navier-Stokes equation is solved through the lattice-Boltzmann method (LBM), see [36] for more details. Our LBM solver uses the D3Q19 lattice, Guo’s forcing scheme [40], the Bhatnagar-Gross-Krook (BGK) collision operator [41], the Bouzidi-Firdaouss-Lallemand no-slip boundary condition at the vessel walls [42], and the Ladd velocity boundary condition for open boundary velocity boundary conditions [43]. Table S1 shows the parameters to solve the LBM in this work.

**Table S1.**
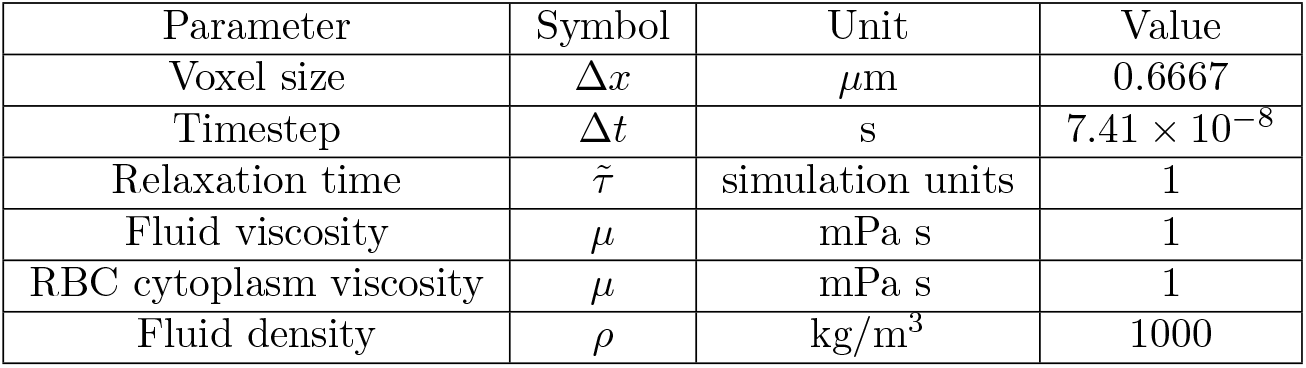
Simulation parameters used for the LBM. Parameters with a tilde are in simulation units.

| Parameter | Symbol | Unit | Value |
| --- | --- | --- | --- |
| Voxel size | $\Delta x$ | $\mu\text{m}$ | 0.6667 |
| Timestep | $\Delta t$ | s | $7.41 \times 10^{-8}$ |
| Relaxation time | $\tilde{\tau}$ | simulation units | 1 |
| Fluid viscosity | $\mu$ | mPa s | 1 |
| RBC cytoplasm viscosity | $\mu$ | mPa s | 1 |
| Fluid density | $\rho$ | kg/m<sup>3</sup> | 1000 |

### Red blood cell model

The red blood cell (RBC) membrane is treated as an isotropic and hyperelastic two-dimensional surface having the shape of a biconcave disk, similarly to a physiological RBC [44]. Numerically, the membrane is discretised into *N*_*f*_ triangular faces *j*, onto which the equations for the membrane model are discretised. Below are given the discretised equations for the membrane model. The RBC membrane is modelled with a membrane energy

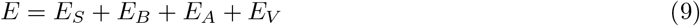

where the subscripts S, B, A, V denote the energy contributions from strain, bending, area and volume energies, respectively. Our model uses the strain energy density from Skalak et al. [45],

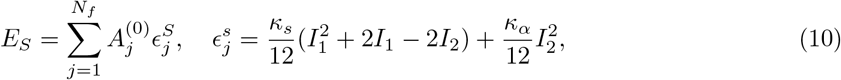

where *κ*_*s*_ and *κ*_*α*_ are the shear and dilation moduli, *A* is the surface area of a membrane triangle and the superscript (0) denotes the underformed state. *I*_1_ and *I*_2_ are the strain invariants, see [46] for details on how to calculate them.

The bending energy is calculated from [47]

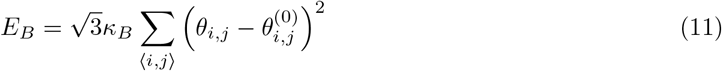

where *κ*_*β*_ is the bending modulus and *θ*_*i,j*_ is the angle between two neighbouring triangular faces.

In addition, the area and volume energy contributions, *E*_*A*_ and *E*_*V*_, impose a penalty on deviation of the RBC membrane from its rest surface area and volume, thus enforcing a quasi constant surface area and volume for the RBCs.

The forces acting on each node of the RBC mesh can then be calculated through the concept of virtual work,

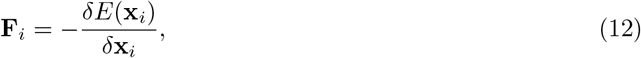

where **F**_*i*_ is the force acting on the *i*^*th*^ node of the membrane mesh and **x**_*i*_ is the position of the node. Table S2 shows the parameters for the RBC model.

**Table S2.**
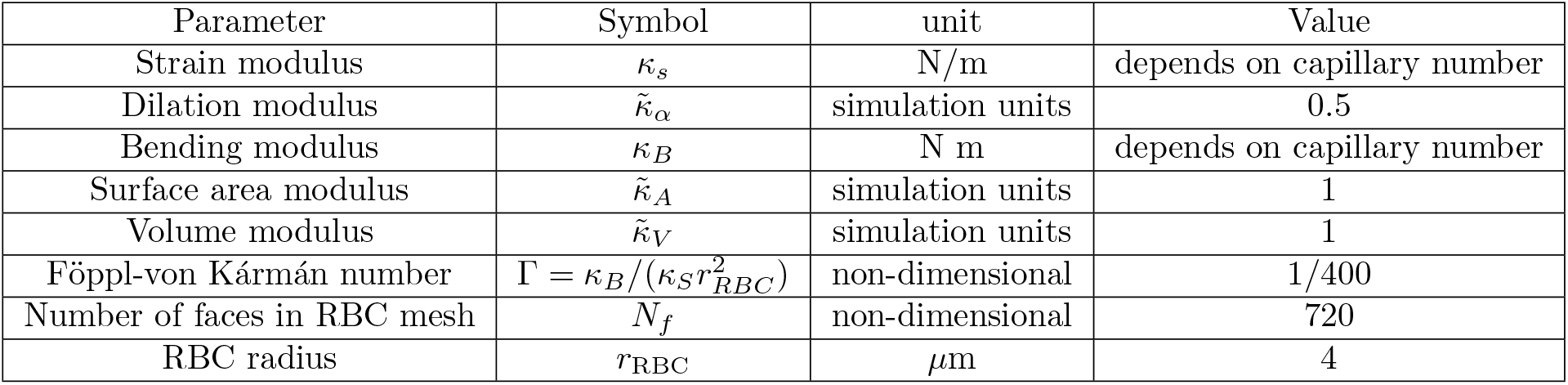
Parameters used for the RBC model. Parameters with a tilde are in simulation units.

### Fluid-cell interaction

The RBC membrane is bi-directionally coupled to the suspending fluid, where the no-slip condition between the RBC membrane and the fluid is imposed and the forces resisting deformation of the membrane also act on the suspending fluid. The bi-directional coupling is numerically modelled through the immersed boundary method (IBM) [35, 48].

The initial step consists in spreading the forces acting on the membrane nodes, from Eq. (12), to the fluid through

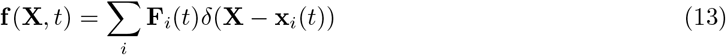

where **f** (**X**, *t*) is the force density acting on the fluid node at position **X** and time *t*, and *δ*(**X − x**_*i*_(*t*)) is the discretised delta function for which we use a three-point stencil [35]. **f** (**X**, *t*) is imposed onto the fluid as an external force in the LBM.

The no-slip condition is enforced through an interpolation of the fluid velocity at the locations of each RBC membrane node through

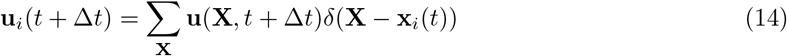

where **u**_*i*_(*t* + Δ*t*) is the velocity of the *i*^*th*^ RBC mesh node at time *t* + Δ*t* and **u**(**X**, *t* + Δ*t*) is the updated velocity at a fluid lattice point **X**. The same three-point stencil is used. Finally, the nodes on the RBC mesh are advected through

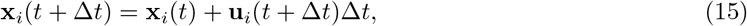

enforcing the no-slip boundary condition as the membrane nodes move at the same velocity as the suspending fluid.

### Network blood flow model

#### Poiseuille model

In this model, blood in an individual vessel segment is treated as a continuous and incompressible fluid using Poiseuille’s law, assuming that blood has an apparent viscosity. Poiseuille’s law can be expressed, for a circular and elliptical cross section, respectively:

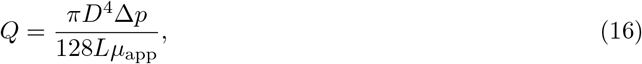

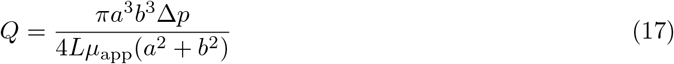

where *Q* is the fluid flow rate, *D* is the vessel diameter, Δ*p* is the pressure drop along the vessel segment of length *L*, and *μ*_app_ is the apparent viscosity of the fluid. In the case of an elliptical cross-section, *a* and *b* are the major and minor radii of the ellipse, and one recovers Poiseuille’s law for a circular cross-section for *a* = *b*.

The apparent viscosity of blood is treated through an empirically obtained function relating the apparent viscosity to the vessel diameter and the discharge haematocrit in the vessel:

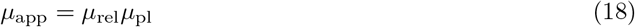

where *μ*_pl_ is the viscosity of plasma. Furthermore, the relative viscosity *μ*_rel_ can be calculated through

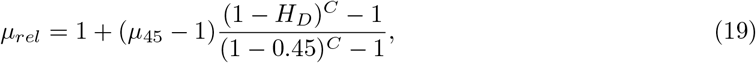

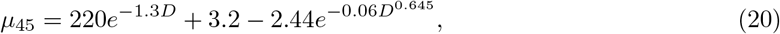

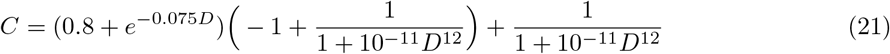

where *H*_*D*_ is the discharge haematocrit in the vessel and *D* is the vessel diameter in microns.

With a known discharge haematocrit, one can apply Poiseuille’s law to every vessel segment in a network of vessels, leading to a system of equations where the pressure values at the end of every vessel segment is unknown [19]. By applying boundary conditions at the open-ended vessel segments, the system of equations can be solved, calculating the pressure in the network, therefore allowing one to calculate the flow rate in each vessel segment.

However, the discharge haematocrit is not known a priori in each vessel segment, due to the disproportional partitioning of RBCs at diverging bifurcations. The partitioning of RBCs at diverging vascular bifurcations is quantified using Pries’ plasma skimming law [24, 49],

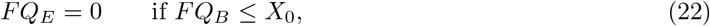

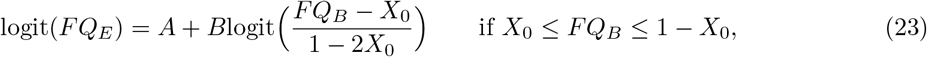

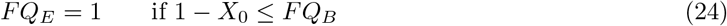

where *FQ*_*e*_ is the fractional flow rate of RBCs to the child branch and *FQ*_*b*_ is the fractional flow rate of blood to the same child branch. The fractional flow rate to a child branch is the flow rate in the said child branch divided by the flow rate of the parent branch. The remaining parameters, *A, B*, and *X*_0_ are calculated through the following equations:

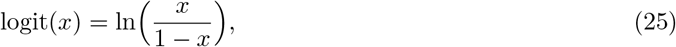

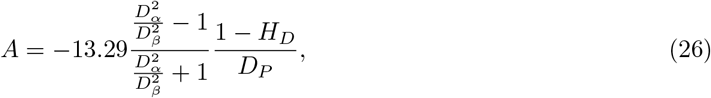

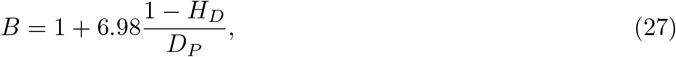

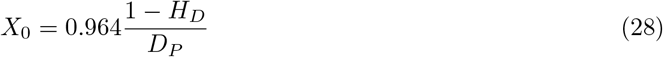

where *D*_*α*_ and *D*_*β*_ are the diameters of the two child branches in microns, *D*_*P*_ is the diameter of the parent branch in microns, and *H*_*D*_ is the discharge haematocrit of the parent branch. With eq. (22)–(24) one can calculate the discharge haematocrit in both child branches.

#### Numerical solution

The aforementioned model is solved through an iterative scheme, as in [19, 50], due to the heterogeneity in haematocrit introduced by the partitioning of RBCs in eq. (22)–(24). At each iteration, a linear system of equations containing Poiseuille’s law at every bifurcation is set up and solved with the known boundary conditions. The haematocrit in the network is then calculated using eq. (22)–(24). The solution of flow rates for blood and RBCs is then compared to the previous iteration solution. The convergence criterion is

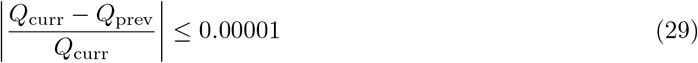

where *Q*_curr_ is the flow rate in a vessel segment for the current iteration and *Q*_prev_ is the flow rate in the same vessel segment for the previous iteration. This equation is applied at every vessel segment, for both blood flow and RBC flow. At the end of an iteration, if the convergence criteria is met for every vessel in the network, the simulation is complete and terminated.

#### Radial and random compression models

**Figure S1.**
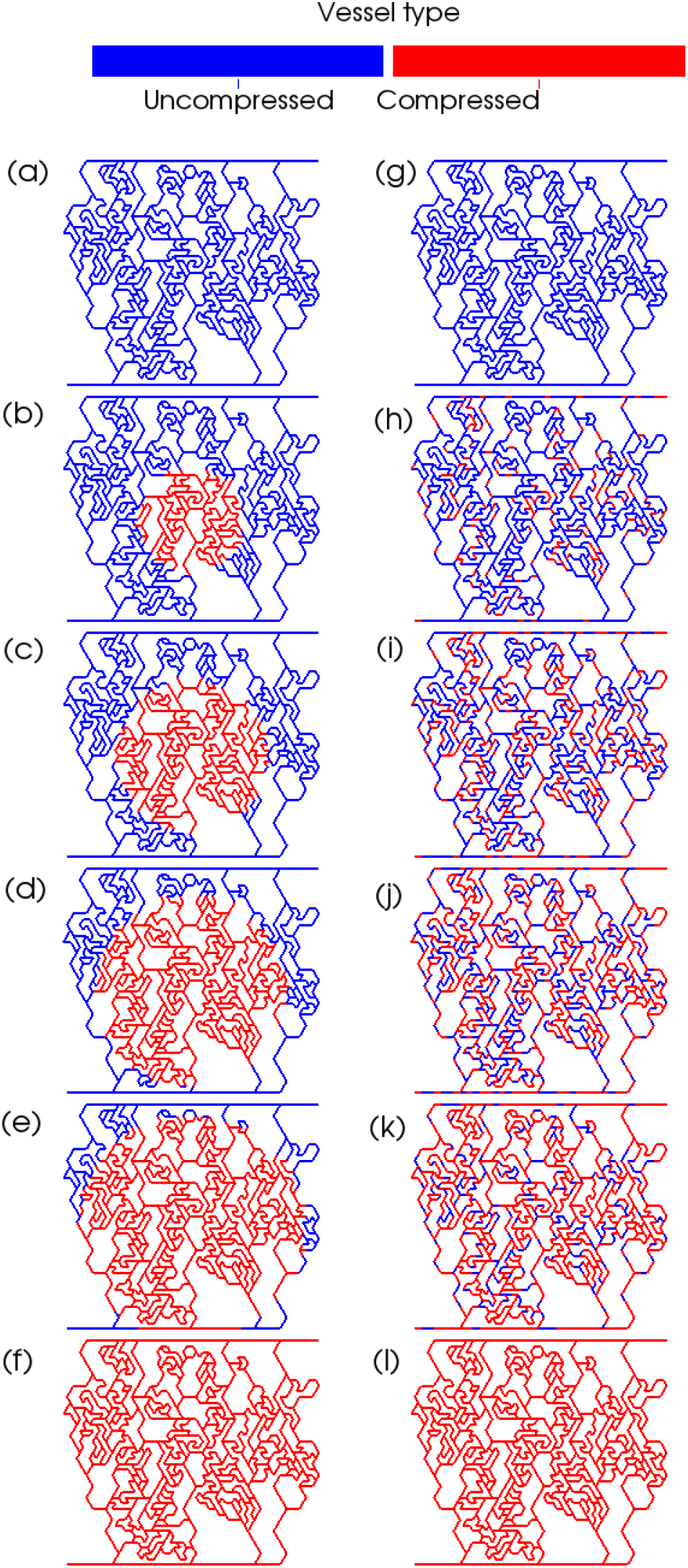
Illustration of network compression models. (a)–(f) illustrate the radial model from 0% to 100% of compressed vessels in steps of 20%. (g)–(l) illustrate the random model from 0% to 100% of compressed vessels in steps of 20%.

#### Critical bifurcation illustration

**Figure S2.**
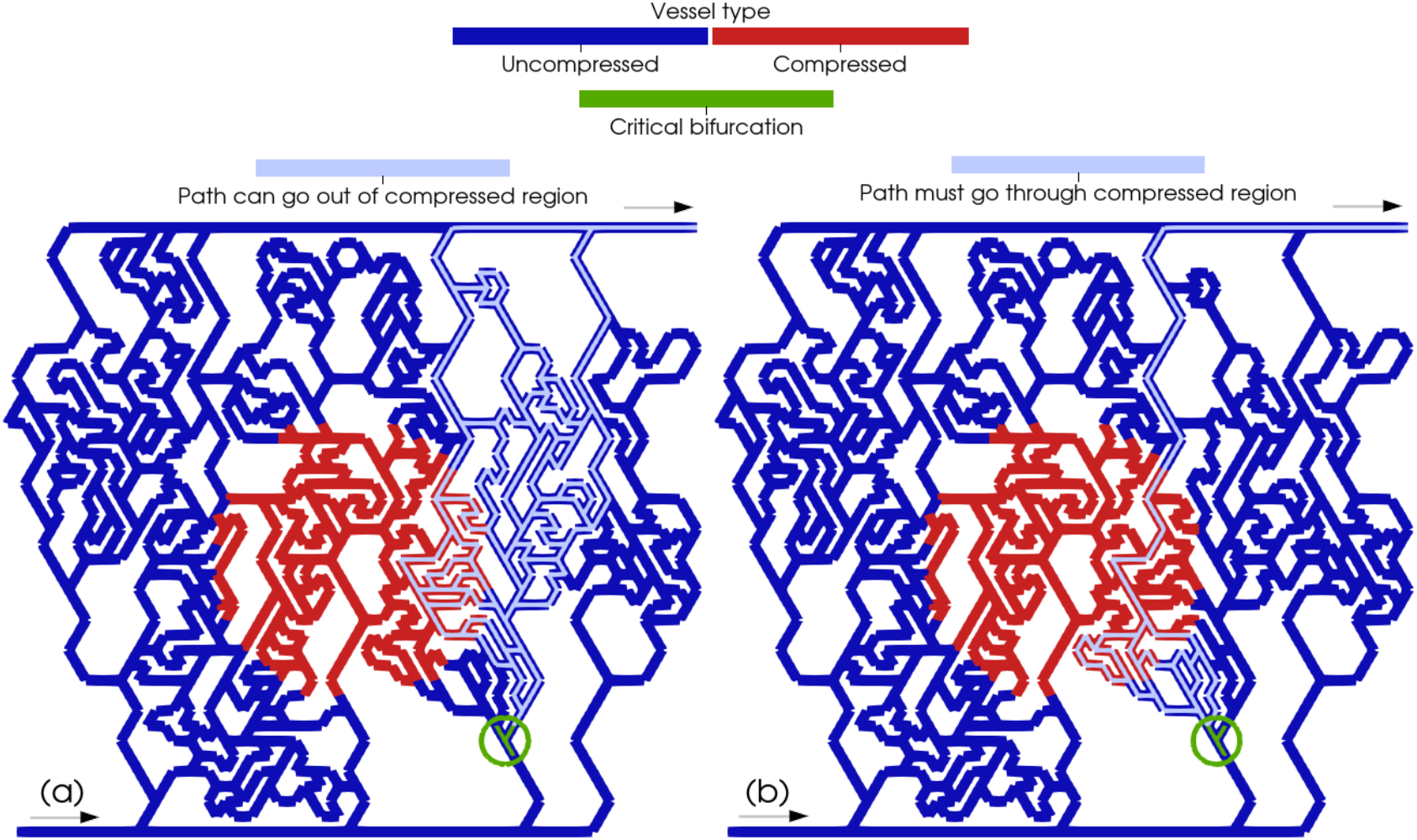
Illustration of one critical bifurcation. In dark blue are the vessels which are treated as non-compressed, and in red the vessels treated as compressed. The green vessels are the vessels in the identified critical bifurcation and are circled in green for visibility. The arrows at the inlet and outlet indicate the direction of flow. (a) Vessels in light blue are all the paths from one of the child branches of the critical bifurcation where at least one path can go around the compressed region. (b) Vessels in light blue are all the paths from one of the child branches of the critical bifurcation where all paths must go through the compressed region.

## Table of simulations to parameterise model

**Table S3.**
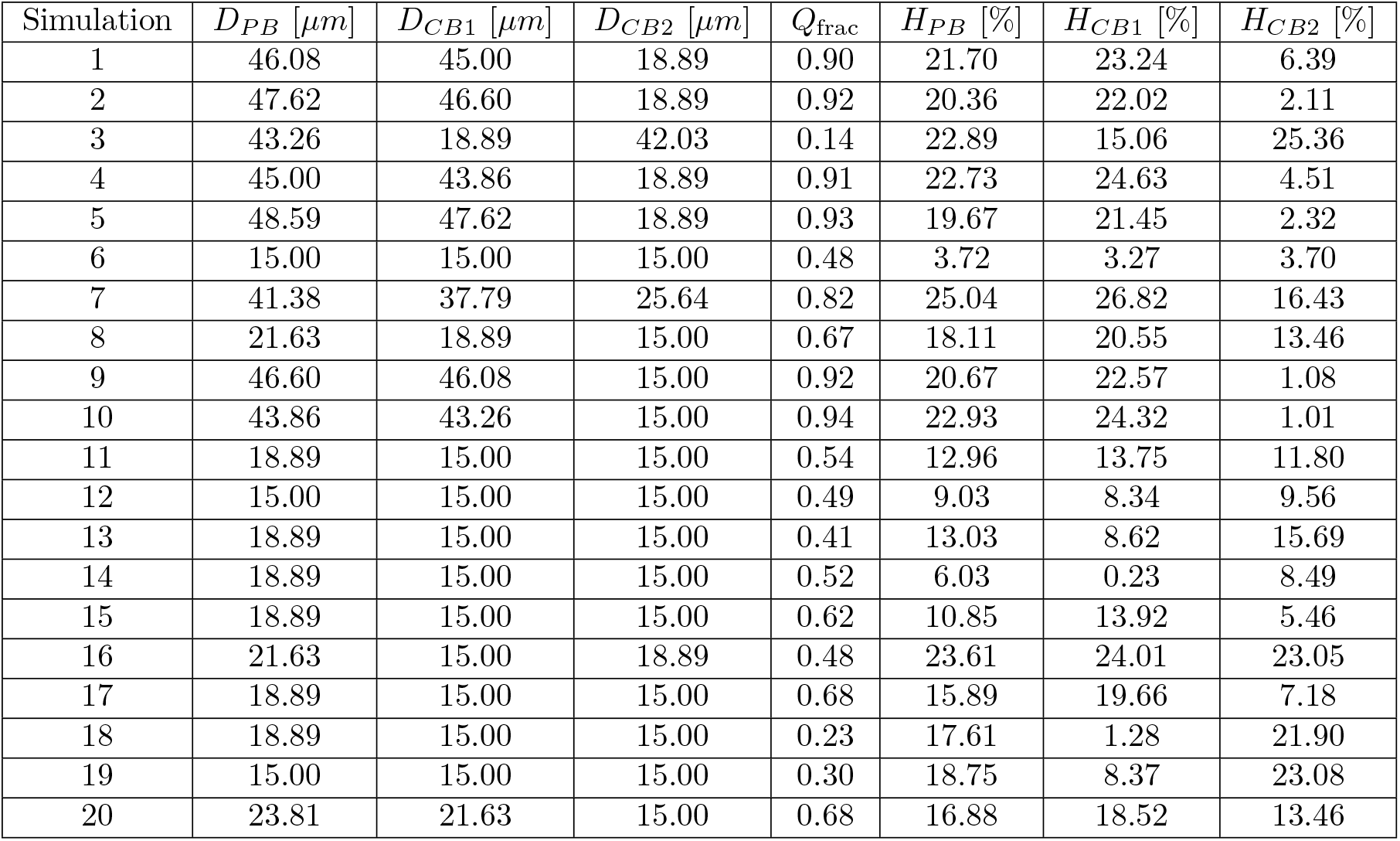
Parameters of the fully resolved simulations to calculate an updated *X*_0_ parameter for compressed vessels. All parent branches are compressed with an aspect ratio of 4.26 [8] as per [13]. *D*_*P B*_ is the undeformed parent branch diameter, *D*_*CB*1_ is the diameter of the first child branch, *D*_*CB*2_ is the diameter of the second child branch, *Q*_frac_ is the flow rate fraction to the first child branch, *H*_*P B*_ is the parent branch haematocrit, *H*_*CB*1_ is the haematocrit of the first child branch, and *H*_*CB*2_ is the haematocrit of the second child branch.

### Derivation for Δ*H*_1_, Δ*H*_2_, **and** 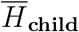

At individual bifurcations where the higher flowing child branch is disproportionately enriched in RBCs, it is possible to show mathematically, with some assumptions, that the average haematocrit of the child branches is lower than the haematocrit of the parent branch. One can start with a mass balance on the blood and RBC flows, using Fig. S3 as a sketch for the system:

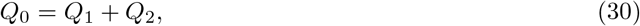

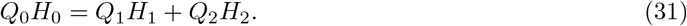

One can then express the haematocrit in the child branches as a deviation from the parent branch haematocrit where one assumes branch 1 to be the enriched branch:

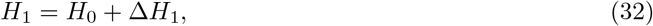

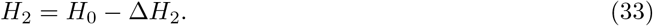

Combining eq. (30) with eq. (31) and substituting in eq. (32) and (33), one can simplify the result to

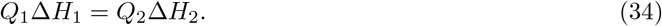

**Figure S3.**
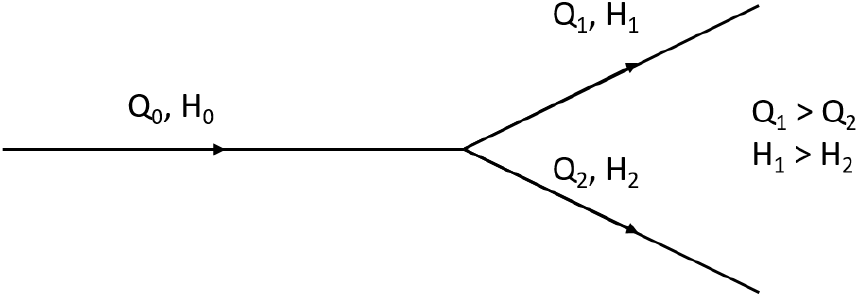
Model bifurcation. *H* is haematocrit, *Q* is flow rate. Subscripts 0, 1, and 2 denote the parent branch and two child branches, respectively. Child branch 1 is higher in flow rate, and therefore in haematocrit.

If one assumes unequal partitioning of blood flow, such that *Q*_1_ > *Q*_2_, then to satisfy equation (34) it must follow that Δ*H*_2_ > Δ*H*_1_.

One may then calculate the average haematocrit of the child branches:

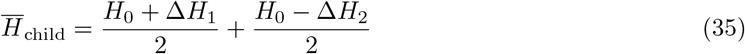

which simplifies to

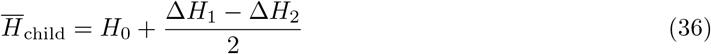

and, with Δ*H*_2_ > Δ*H*_1_, then it follows that

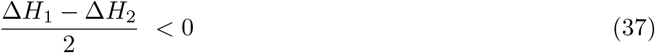

and therefore that

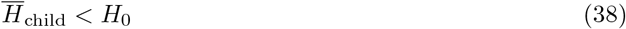

as the fraction on the RHS of the inequality in eq. (36) must be negative, to satisfy eq. (37).

## References

1. Jain, R. K., Martin, J. D. & Stylianopoulos, T. The Role of Mechanical Forces in Tumor Growth and Therapy. Annual Review of Biomedical Engineering 16, 321–346 (2014).

2. Brizel, D. M., Scully, S. P., Harrelson, J. M., Layfleld, L. J., Bean, J. M., Prosnitz, L. R. & Dewhirst, M. W. Tumor Oxygenation Predicts for the Likelihood of Distant Metastases in Human Soft Tissue Sarcoma1 tech. rep. (1996), 941–943.

3. Brown, J. M. & Wilson, W. R. Exploiting tumour hypoxia in cancer treatment. Nature Reviews Cancer 4, 437–447 (2004).

4. Jain, R. K. Normalizing tumor microenvironment to treat cancer: Bench to bedside to biomarkers. Journal of Clinical Oncology 31, 2205–2218 (2013).

5. Martin, J. D., Cabral, H., Stylianopoulos, T. & Jain, R. K. Improving cancer immunotherapy using nanomedicines: progress, opportunities and challenges. Nature Reviews Clinical Oncology 17, 251–266 (Apr. 2020).

6. Munn, L. L. & Jain, R. K. Vascular regulation of antitumor immunity. Science 365, 544–545 (2019).

7. Padera, T., Stoll, B., Tooredman, J., Capen, D., di Tomaso, E. & Jain, R. Cancer cells compress intratumour vessels. Nature 427, 695 (2004).

8. Fang, L., He, Y., Tong, Y., Hu, L., Xin, W., Liu, Y., Zhong, L., Zhang, Y. & Huang, P. Flattened microvessel independently predicts poor prognosis of patients with non-small cell lung cancer. Oncotarget 8, 30092–30099 (2017).

9. Hagendoorn, J., Tong, R., Fukumura, D., Lin, Q., Lobo, J., Padera, T. P., Xu, L., Kucherlapati, R. & Jain, R. K. Onset of abnormal blood and lymphatic vessel function and interstitial hypertension in early stages of carcinogenesis. Cancer Research 66, 3360–3364 (2006).

10. Chauhan, V. P., Martin, J. D., Liu, H., Lacorre, D. A., Jain, S. R., Kozin, S. V., Stylianopoulos, T., Mousa, A. S., Han, X., Adstamongkonkul, P., Popović, Z., Huang, P., Bawendi, M. G., Boucher, Y. & Jain, R. K. Angiotensin inhibition enhances drug delivery and potentiates chemotherapy by decompressing tumour blood vessels. Nature Communications 4 (2013).

11. Secomb, T. W. Blood Flow in the Microcirculation. Annual Review of Fluid Mechanics 49, 443–461 (2017).

12. Bernabeu, M. O., Köry, J., Grogan, J. A., Markelc, B., Beardo, A., D’Avezac, M., Enjalbert, R., Kaeppler, J., Daly, N., Hetherington, J., Krüger, T., Maini, P. K., Pitt-Francis, J. M., Muschel, R. J., Alarcón, T. & Byrne, H. M. Abnormal morphology biases haematocrit distribution in tumour vasculature and contributes to heterogeneity in tissue oxygenation. Proceedings of the National Academy of Sciences 117, 27811–27819 (2020).

13. Enjalbert, R., Hardman, D., Krüger, T. & Bernabeu, M. O. Compressed vessels bias red blood cell partitioning at bifurcations in a hematocrit-dependent manner: Implications in tumor blood flow. Proceedings of the National Academy of Sciences 118 (2021).

14. Goldman, D. Theoretical models of microvascular oxygen transport to tissue. Microcirculation 15, 795–811 (2008).

15. Jain, R. K. Determinants of Tumor Blood Flow: A Review. CANCER RESEARCH 48, 2641–2658 (1988).

16. Seano, G., Nia, H. T., Emblem, K. E., Datta, M., Ren, J., Krishnan, S., Kloepper, J., Pinho, M. C., Ho, W. W., Ghosh, M., Askoxylakis, V., Ferraro, G. B., Riedemann, L., Gerstner, E. R., Batchelor, T. T., Wen, P. Y., Lin, N. U., Grodzinsky, A. J., Fukumura, D., Huang, P., Baish, J. W., Padera, T. P., Munn, L. L. & Jain, R. K. Solid stress in brain tumours causes neuronal loss and neurological dysfunction and can be reversed by lithium. Nature Biomedical Engineering 3, 230–245 (Mar. 2019).

17. Varchanis, S., Dimakopoulos, Y., Wagner, C. & Tsamopoulos, J. How viscoelastic is human blood plasma? Soft Matter 14, 4238–4251 (2018).

18. Mazzeo, M. D. & Coveney, P. V. HemeLB: A high performance parallel lattice-Boltzmann code for large scale fluid flow in complex geometries. Computer Physics Communications 178, 894–914 (2008).

19. Pries, A. R., Secomb, T. W., Gaehtgens, P. & Gross, J. F. Blood flow in microvascular networks. Experiments and simulation. Circulation research, 826–834 (1990).

20. Rieger, H. & Welter, M. Integrative models of vascular remodeling during tumor growth May 2015.

21. Fredrich, T., Welter, M. & Rieger, H. Tumorcode: A framework to simulate vascularized tumors. European Physical Journal E 41 (Apr. 2018).

22. Rieger, H., Fredrich, T. & Welter, M. Physics of the tumor vasculature: Theory and experiment Feb. 2016.

23. Stylianopoulos, T., Martin, J. D., Snuderl, M., Mpekris, F., Jain, S. R. & Jain, R. K. Coevolution of solid stress and interstitial fluid pressure in tumors during progression: Implications for vascular collapse. Cancer Research 73, 3833–3841 (2013).

24. Pries, A., Ley, K., Classen, M. & Gaehtgens, P. Red Cell Distribution at Microvascular. Microvascular research 38, 81–101 (1989).

25. Pries, A. R., Ley, K. & Gaehtgens, P. Generalization of the Fahraeus principle for microvessel networks. American Journal of Physiology - Heart and Circulatory Physiology 251 (1986).

26. Nia, H. T., Munn, L. L. & Jain, R. K. Physical traits of cancer. Science 370 (2020).

27. Kamoun, W. S., Chae, S. S., Lacorre, D. A., Tyrrell, J. A., Mitre, M., Gillissen, M. A., Fukumura, D., Jain, R. K. & Munn, L. L. Simultaneous measurement of RBC velocity, flux, hematocrit and shear rate in vascular networks. Nature Methods 7, 655–660 (2010).

28. Merlo, A., Berg, M., Duru, P., Risso, F., Davit, Y. & Lorthois, S. A few upstream bifurcations drive the spatial distribution of red blood cells in model microfluidic networks. Soft Matter 18, 1463–1478 (Feb. 2022).

29. Sweeney, P. W., D’Esposito, A., Walker-Samuel, S. & Shipley, R. J. Modelling the transport of fluid through heterogeneous, whole tumours in silico. PLoS computational biology 15, e1006751 (2019).

30. d’Esposito, A., Sweeney, P. W., Ali, M., Saleh, M., Ramasawmy, R., Roberts, T. A., Agliardi, G., Desjardins, A., Lythgoe, M. F., Pedley, R. B., Shipley, R. & Walker-Samuel, S. Computational fluid dynamics with imaging of cleared tissue and of in vivo perfusion predicts drug uptake and treatment responses in tumours. Nature Biomedical Engineering 2, 773–787 (2018).

31. Welter, M., Fredrich, T., Rinneberg, H. & Rieger, H. Computational model for tumor oxygenation applied to clinical data on breast tumor hemoglobin concentrations suggests vascular dilatation and compression. PLoS ONE 11, 1–42 (2016).

32. Dewhirst, M. W., Mowery, Y. M., Mitchell, J. B., Cherukuri, M. K. & Secomb, T. W. Rationale for hypoxia assessment and amelioration for precision therapy and immunotherapy studies. Journal of Clinical Investigation 129, 489–491 (Jan. 2019).

33. Dewhirst, M. W. & Secomb, T. W. Transport of drugs from blood vessels to tumour tissue. Nature Reviews Cancer 17, 738–750 (2017).

34. Jain, R. K. Normalization of Tumor Vasculature: An Emerging Concept in Antiangiogenic Therapy. Science 307, 58–62 (2005).

35. Krüger, T., Varnik, F. & Raabe, D. Efficient and accurate simulations of deformable particles immersed in a fluid using a combined immersed boundary lattice Boltzmann finite element method. Computers and Mathematics with Applications 61, 3485–3505 (2011).

36. Kruger, T., Kusumaatmaja, H., Kuzmin, A., Shardt, O., Goncalo, S. & Viggen, E. M. The lattice boltzmann method, principles and practice 207, 1–705 (Springer, 2017).

37. Krueger, T. margination and near-wall dynamics. Rheologica Acta (2015).

38. Krüger, T., Holmes, D. & Coveney, P. V. Deformability-based red blood cell separation in deterministic lateral displacement devices-A simulation study. Biomicrofluidics 8 (Oct. 2014).

39. Zhou, Q., Fidalgo, J., Bernabeu, M. O., Oliveira, M. S. N. & Krüger, T. Emergent cell-free layer asymmetry and biased haematocrit partition in a biomimetic vascular network of successive bifurcations. Soft Matter (2021).

40. Guo, Z., Zheng, C. & Shi, B. Discrete lattice effects on the forcing term in the lattice Boltzmann method. Physical Review E -Statistical Physics, Plasmas, Fluids, and Related Interdisciplinary Topics 65, 1–6 (2002).

41. Bhatnagar, P. L., Gross, E. P. & Krook, M. A Model for Collision Processes in Gases. I. Small Amplitude Processes in Charged and Neutral One-Component Systems. Phys. Rev. 94 (1954).

42. Bouzidi, M., Firdaouss, M. & Lallemand, P. Momentum transfer of a Boltzmann-lattice fluid with boundaries. Physics of Fluids 13, 3452–3459 (2001).

43. Ladd, A. J. Numerical Simulations of Particulate Suspensions Via a Discretized Boltzmann Equation. Part 1. Theoretical Foundation. Journal of Fluid Mechanics 271, 285–309 (1994).

44. Evans, E. & Fung, Y.-C. Improved Measurements of the Erythrocyte Geometry. Microvascular research 4, 335–347 (1972).

45. Skalak, R., Tozeren, A., Zarda, R. P. & Chien, S. Strain Energy Function of Red Blood Cell Membranes. Biophysical Journal 13, 245–264 (1973).

46. Timm Krüger. Computer simulation study of collective phenomena in dense suspensions of red blood cells under shear PhD thesis (2012).

47. Guckenberger, A. & Gekle, S. Theory and algorithms to compute Helfrich bending forces: A review Apr. 2017.

48. Peskin, C. S. The immersed boundary method. Acta Numerica 11, 479–517 (Jan. 2002).

49. Pries, A. R. & Secomb, T. W. Microvascular blood viscosity in vivo and the endothelial surface layer. American Journal of Physiology - Heart and Circulatory Physiology 289, 2657–2664 (2005).

50. Lorthois, S., Cassot, F. & Lauwers, F. Simulation study of brain blood flow regulation by intracortical arterioles in an anatomically accurate large human vascular network: Part I: Methodology and baseline flow. NeuroImage 54, 1031–1042 (Jan. 2011).

